# The emergence of SARS-CoV-2 in Europe and the US

**DOI:** 10.1101/2020.05.21.109322

**Authors:** Michael Worobey, Jonathan Pekar, Brendan B. Larsen, Martha I. Nelson, Verity Hill, Jeffrey B. Joy, Andrew Rambaut, Marc A. Suchard, Joel O. Wertheim, Philippe Lemey

## Abstract

Accurate understanding of the global spread of emerging viruses is critically important for public health response and for anticipating and preventing future outbreaks. Here, we elucidate when, where and how the earliest sustained SARS-CoV-2 transmission networks became established in Europe and the United States (US). Our results refute prior findings erroneously linking cases in January 2020 with outbreaks that occurred weeks later. Instead, rapid interventions successfully prevented onward transmission of those early cases in Germany and Washington State. Other, later introductions of the virus from China to both Italy and Washington State founded the earliest sustained European and US transmission networks. Our analyses reveal an extended period of missed opportunity when intensive testing and contact tracing could have prevented SARS-CoV-2 from becoming established in the US and Europe.

## Introduction

The emergence in late 2019 of SARS-CoV-2, the agent of COVID-19, has ignited a global health crisis unparalleled since the 1918 influenza pandemic. Rapid sharing of viral sequence data that began within weeks of identification of the virus (*1*) allowed the development of diagnostic tests crucial to control efforts and initiated research enabling candidate vaccines and therapies. These genomes also precipitated a worldwide effort of viral genomic sequencing and analyses unprecedented in scale and pace. At time of writing (May 15^th^, 2020) there are already 25,181 complete genomes available.

Embedded within this assemblage of genomic data is crucial information about the nature and history of the pandemic. Here, we investigate fundamental questions about when, where and how this new virus established itself globally—scientific questions that have become inextricably linked to important societal and policy concerns. The combination of the relatively slow rate of SARS-CoV-2 evolution, its rapid dissemination within and between locations, and extremely unrepresentative sampling means that naive interpretation of phylogenetic analyses can lead to serious error. However, we show that with the integration of multiple sources of information, careful consideration of evolutionary process and pattern, and cutting-edge technology, key events in the unfolding of the pandemic, which have been the subject of conjecture and controversy but little quantitative study, can be resolved.

Here we consider two of the earliest known introductions—successful or not—of SARS-CoV-2 into both Europe and the US.

The first patient to be diagnosed with COVID-19 in the US, designated ‘WA1’, was a Chinese national who travelled from Wuhan, in Hubei Province, China, to Sea-Tac International Airport near Seattle, Washington State, arriving on January 15^th^, 2020 (*2*). This individual was attuned to Centers for Disease Control and Prevention (CDC) messaging about the new pneumonial disease circulating in Wuhan and promptly sought medical care upon becoming symptomatic with COVID-19, receiving a sequence-confirmed diagnosis and becoming the first US patient to have a SARS-CoV-2 genome sequenced, sampled on January 19^th^, 2020 (*3*). Efficient contact tracing measures were enacted by local health authorities and the patient and public health authorities worked together in an exemplary fashion to limit opportunities for spread (*3*). No onward transmission was detected after exhaustive follow-up in what appeared to be successful containment of the first known incursion of the virus in the US (*3*).

On February 29^th^, 2020, a SARS-CoV-2 genome was reported (*4*) from a second Washington State patient, ‘WA2’, whose virus had been sampled on February 24th as part of a community surveillance study of respiratory viruses (*5*). The report’s authors calculated a high probability that WA2 was a direct descendent of WA1, coming to the surprising conclusion that there had by that point already been six weeks of cryptic circulation of the virus in Washington State (*4*). The finding, described in a lengthy Twitter thread on February 29th, fundamentally altered the picture of the SARS-CoV-2 situation in the US, and seemed to show how the power of genomic epidemiology could be harnessed to uncover hidden epidemic dynamics and inform policy making in real time (*6*, *7*). The COVID-19 pandemic represents the first major global disease event to emerge in the age of social media, and the urgent need for timely information to inform public health decisions has frequently outpaced standard peer-review processes. As a result, the findings played a decisive role in Washington State’s early adoption of intensive social distancing efforts, which, in turn, appeared to explain Washington State’s relative success in controlling the outbreak, compared with states that delayed, such as New York (*8*).

In Europe, an employee of the auto supplier Webasto visited the company’s headquarters in Bavaria, Germany, from Shanghai, China, on January 20^th^, 2020 (*9*). She had been infected with SARS-CoV-2 in Shanghai (after her parents had visited from Wuhan) (*10*) and transmitted the virus to a German man who tested positive on January 27th (*11*) and whose viral genome (‘BavPat1’) was sampled on January 28^th^ (*10*). All told, the outbreak infected 16 employees but was apparently contained through rapid testing and isolation, in efforts that were lauded as an impressive, early display of how to stop the virus (*9*).

Weeks later, however, Italy’s first major outbreak in Lombardy was associated with viruses closely related to BavPat1, differing by just one nucleotide in the nearly 30,000 nucleotide genome. At a time when social and news media outpaced scientific peer-review, a narrative took hold that the virus from Germany had in fact not been contained but had been transmitting undetected for weeks and had been carried to Italy by an infected German (*9*, *12*). In addition to igniting a devastating outbreak in Italy, this particular lineage (B.1) subsequently spread widely across Europe, seeding outbreaks in many countries (*13*).

Importantly, this lineage also spread to North America, establishing an early beachhead in New York City (*14*, *15*). Furthermore, the B.1 lineage harbors a D614G mutation in the Spike protein that has been controversially claimed to make the virus more contagious (*16*). Greater clarity about the role of that initial outbreak in Germany in the emergence of SARS-CoV-2 in Italy—and then the rest of Europe and the US—has critical implications for individuals involved in the German outbreak and response as well as for the broader feasibility of controlling the virus through gold standard public health responses.

## Results

### Emergence of SARS-CoV-2 in Washington State

The availability of substantially more genomic sequence data, revealing that WA2 belonged to a large, monophyletic ‘WA outbreak’ clade (*17*), affords us the opportunity to revisit whether this outbreak actually began in January with WA1. This narrative has important implications not just for individuals and public health workers in Washington State associated with the WA1 case, but also for evaluating the effectiveness of state- and national-level mitigation strategies in the weeks following the first detection of SARS-CoV-2 in the US.

First, we noted that the ~3% probability reported by Bedford et al (*4*, *17*) that WA1 and the WA outbreak clade would appear so close on the tree by chance is an underestimate: at least four transmission chains co-circulated in the state (their Figure 2), increasing the probability of sampling one that happened to have a close relationship to a rare lineage to >11%. Next, we note a peculiar feature of the relationship between WA1 and the WA outbreak clade. They differ by two substitutions, C17747T and A17858G, and despite hundreds of genomes sequenced in Washington State, no viruses with genomes identical to WA1 or transitional between it and the outbreak clade (i.e. having a C at position 17747 or an A at position 17858, like WA1) (*17*) had been sampled there. This absence was surprising given that scores of genomes identical to the putative descendant of WA1—and inferred most recent common ancestor (MRCA) of the WA outbreak clade—encoding both C17747T and A17858G, have been sampled from Feb 20^th^ until April 27^th^ (Supplementary text). Indeed, a range of phylogenetic patterns could have been observed (Fig. 1A-C), yet were not (Fig. 1D).

**Fig. 1.**
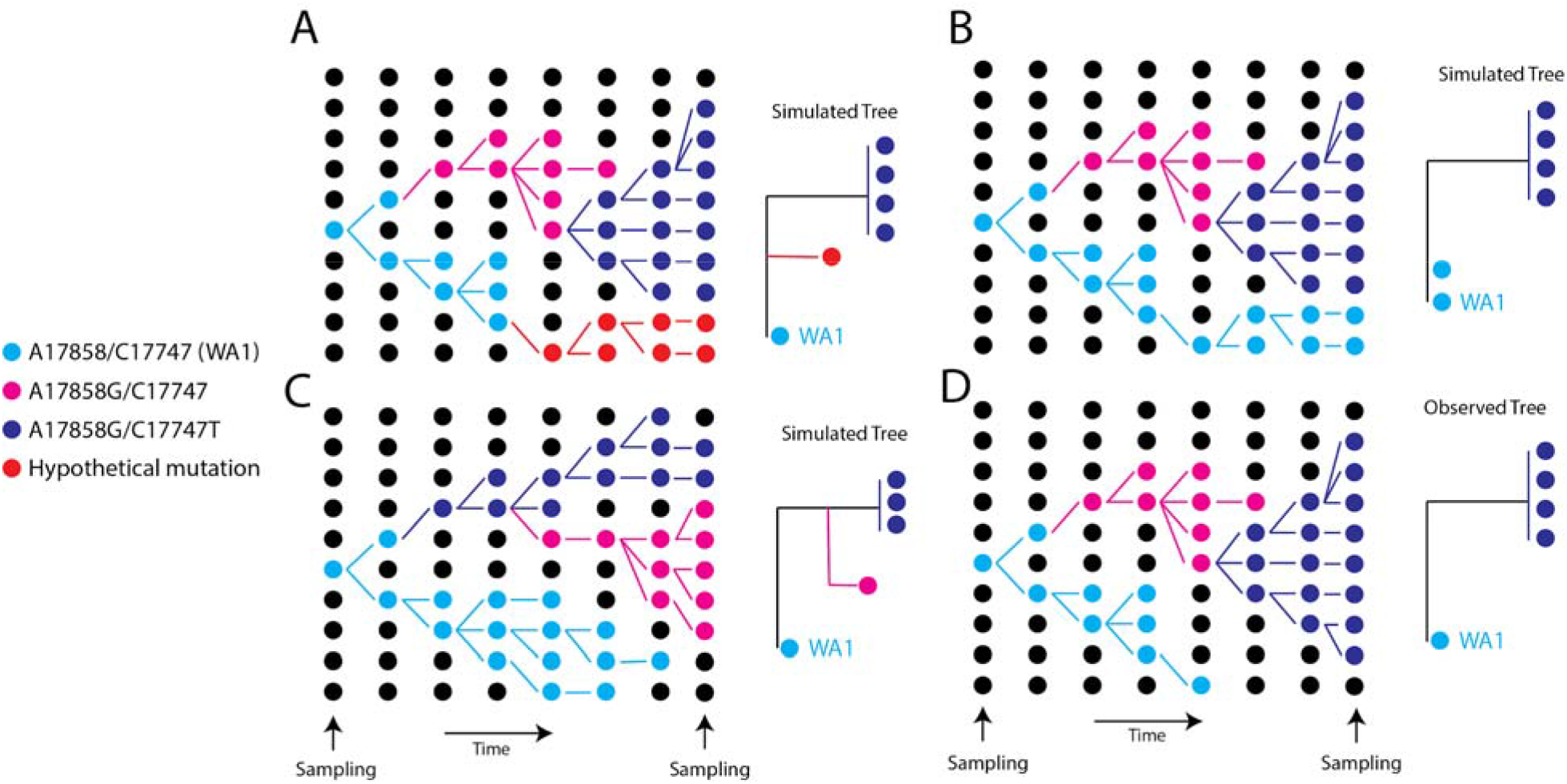
Schematic showing a hypothetical path along which the key mutations in the WA outbreak could have taken in a susceptible population, alongside the inferred phylogeny. (A) Scenario where a hypothetical mutation occurs from WA1-like genomes (B) A hypothetical phylogeny where A17747 and C17858 from the original WA1 virus are maintained in the population and sampled at the end. (C) Hypothetical scenario where a virus one mutation (A17747C) different from WA1 is maintained in the population. (D) The observed tree from the WA outbreak.

**Fig. 2.**
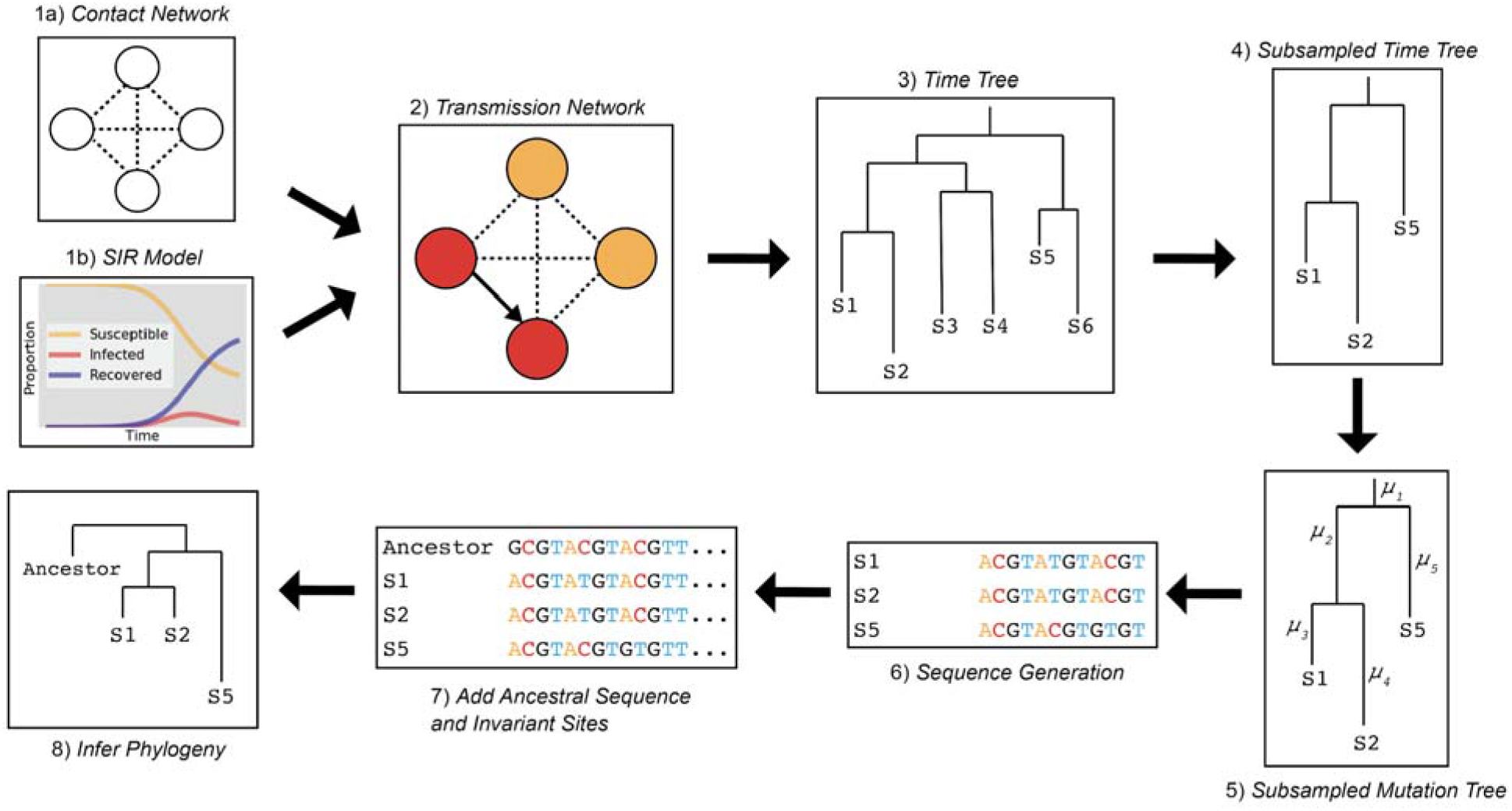
Epidemic simulation workflow. (1) FAVITES generates the contact network and (1a) runs an SIR model (1b) to simulate spread through the contact network and (2) produce a transmission network. (3) FAVITES outputs a viral time tree based on the infected individuals in the transmission network from which (4) individuals are subsampled to match the dates of the original epidemic (e.g., WA outbreak clade). (5) Evolutionary rates are applied to the time tree based on the number of variant sites from the original alignment, converting the branches from years to substitutions/site (μ). (6) Genetic sequences at variant sites are evolved over the subsampled tree using Pyvolve, starting with the ancestral sequence (e.g., WA1), based on GTR parameters inferred from the original alignment. (7) The ancestral sequence and invariant sites are added to the sequence data so (8) a maximum likelihood phylogeny can be inferred in IQ-TREE2.

To investigate whether the observed pattern of evolution was consistent with the WA outbreak clade having descended from WA1, we simulated the entire WA outbreak clade, including the transmission history between thousands of cases, under realistic epidemiologic parameters (*18*), and then evolved genomes forward in time under the constraint that they originated from WA1 (Fig. 2). We simulated 1,000 epidemics seeded by WA1 on January 15^th^ 2020 with a median doubling time of 4.7 days (95% range: 4.2-5.1) and an evolutionary rate of 0.8×10^−3^substitutions/site/year. These simulated epidemics produced a median of 4,269 cases after 61 days (95% range: 1,993-11,053 cases). The median genetic distance from WA1 to the sub-sampled viruses was 3 mutations, which is consistent with the observed phylogeny.

We examined the phylogenetic structure of maximum likelihood trees inferred from sub-sampled simulated viral sequences to determine how frequently they matched the observed relationship between WA1 and the WA outbreak clade. Specifically, a simulation tree matching the observed tree must produce a single branch emanating from WA1 that experiences at least two mutations (C17747T and A17858G in the observed tree) prior to establishment of a single outbreak clade (Fig. 3A). Alternative patterns include: (i) a virus identical to WA1 (Fig. 3B); (ii) a virus that differs from WA1 by a single mutation (Fig. 3C); (iii) a viral lineage forming a basal polytomy with WA1 and the outbreak clade (Fig. 3D); and (iv) a viral lineage that is sibling to the outbreak clade but only experienced a single mutation before divergence (Fig. 3E). The frequency of alternative phylogenetic patterns in the simulated epidemics represents the probability that the true topology (Fig. 3A) could not have occurred if the WA outbreak clade had been seeded by WA1.

**Fig. 3.**
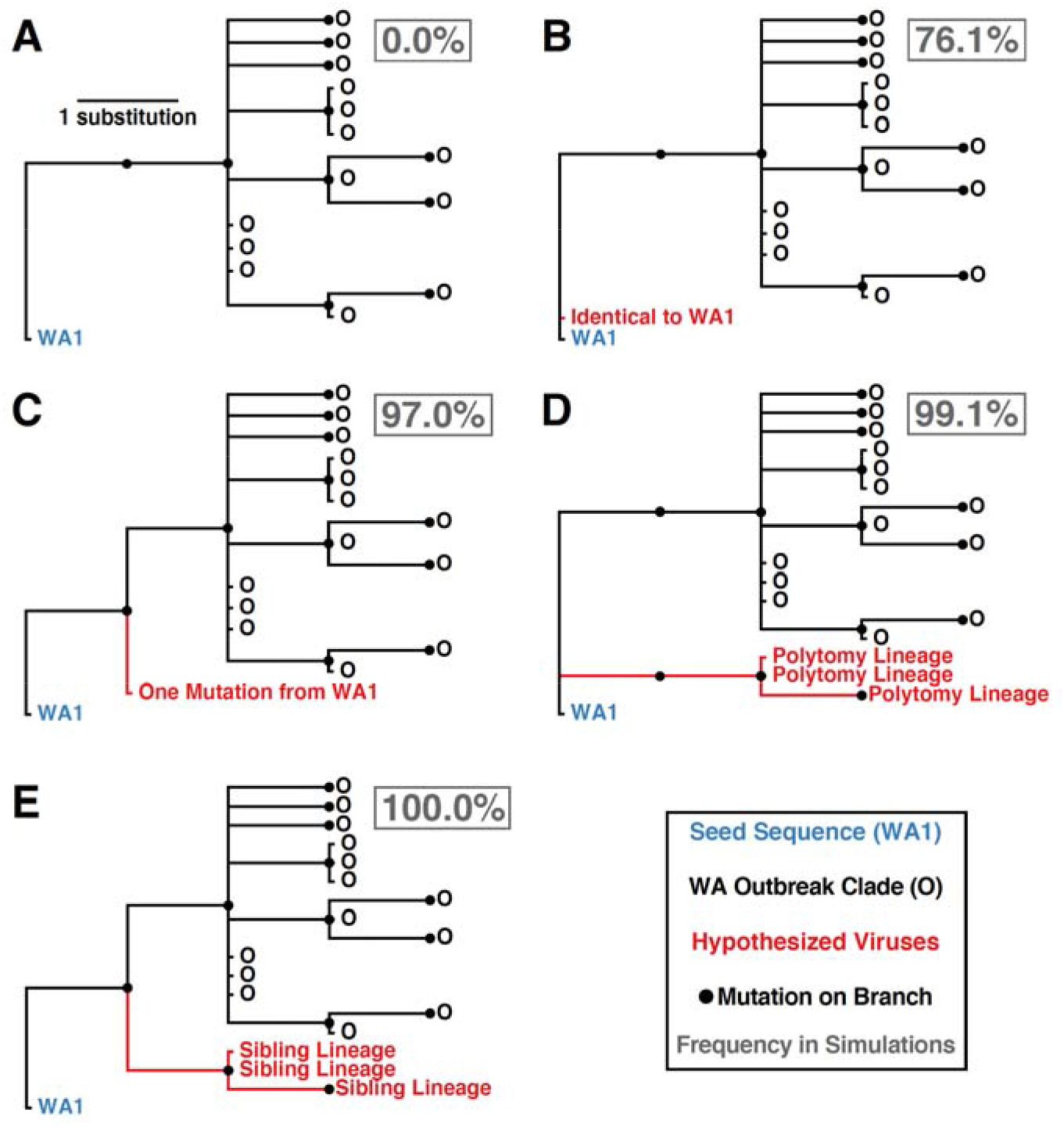
Potential phylogenetic relationships between WA1 and the Washington outbreak clade and their occurrence frequencies in 1000 epidemic simulations. (A) Observed pattern where the WA1 genome is the direct ancestor of the outbreak clade, separated by at least two mutations. (B) Identical sequence to WA1. (C) Sequence that is one mutation divergent from WA1. (D) Lineage forming a basal polytomy with WA1 and the outbreak clade. (E) Sibling lineage to the outbreak clade experiencing only a single mutation from WA1 before divergence. Frequency each relationship was observed in 1000 simulations reported in gray box.

In 76.1% of simulations, we observed at least one virus genetically identical to WA1, with a median of 12 identical viruses in each simulation (95% range: 0-74 identical viruses). Not observing a virus identical to WA1 in the real Washington data does not significantly differ from expectation (*p*=0.239). However, viruses with one mutation from WA1 were observed in 97.0% of simulations, indicating a low probability of not detecting a single sequence from Washington within one mutation of WA1 (*p*=0.030). Lineages forming a basal polytomy with WA1 and the epidemic clade were observed in 99.1% of populations (*p*=0.009) and 100% of simulations had at least one sibling lineage diverging prior to the formation of the outbreak clade (*p*<0.001). Therefore, even if C17747T and A17858G were linked—a possibility since they are both non-synonymous mutations located in the nsp13 helicase gene—we would still expect to see descendants of their predecessors in Washington. In summary, when we seeded the Washington outbreak simulations with WA1 on January 15^th^, 2020, we failed to observe a single simulated epidemic that has the characteristics of the real phylogeny (Fig. 3). These findings are robust to simulations that used a slower epidemic doubling time of 5.6 days (95% range 5.2-5.9) or an accelerated mutation rate of 1.6×10^3^substitutions/site/year (*18*).

#### Accounting for geographical gaps in genomic sampling

A major limitation in phylogeographic inference of SARS-CoV-2 to date has been the low availability of genomic sequence data from locations that experienced early outbreaks, including the original epicenter in Hubei Province, China. Although we do not have access to missing genome sequences, we can estimate how many such genomes are likely missing. We therefore developed methodologies that incorporate unsampled viruses into phylogeographic inferences within a Bayesian framework (*18*). We investigated how tree topologies were affected by the inclusion of unsampled viruses assigned to 13 of the most severely undersampled locations both in China and globally, based on COVID-19 incidence data (*18*). Realistic sampling time distributions also were inferred from incidence data. To better inform placement of unsampled viruses on the phylogeography, we developed a generalized linear model formulation that incorporates air passenger flows between regions within China and between Chinese regions and other countries.

The resulting phylogeny (Fig. 4) provides a reconstruction of the evolutionary relationships of viruses from Washington State that realistically accounts for major gaps in sequence data from Hubei Province, China. For low-diversity data, a single MCC phylogeny has a resolution that is to a large extent not supported by the full posterior tree distribution, but key nodes yield good support in this case including their location estimates. The accommodation of unsampled viruses assigned to Hubei strongly supports independent introductions of WA1 and the WA outbreak clade from Hubei, with ancestral location states supported by high posterior probabilities (≥0.99). Furthermore, using Markov jump estimates that account for phylogenetic uncertainty (*19*), we inferred February 13^th^, 2020 (95% HPD February 7^th^–February 19^th^) as the time, along the branch leading up the MRCA of the WA outbreak clade, at which the founding virus of the WA outbreak clade arrived in Washington State from Hubei. Consistent with these estimates of the introduction date of this viral lineage into Washington State, the Seattle Flu Study tested 6,908 archived samples from January and February, of which only 10 from the end of February were positive (*5*).

**Fig. 4.**
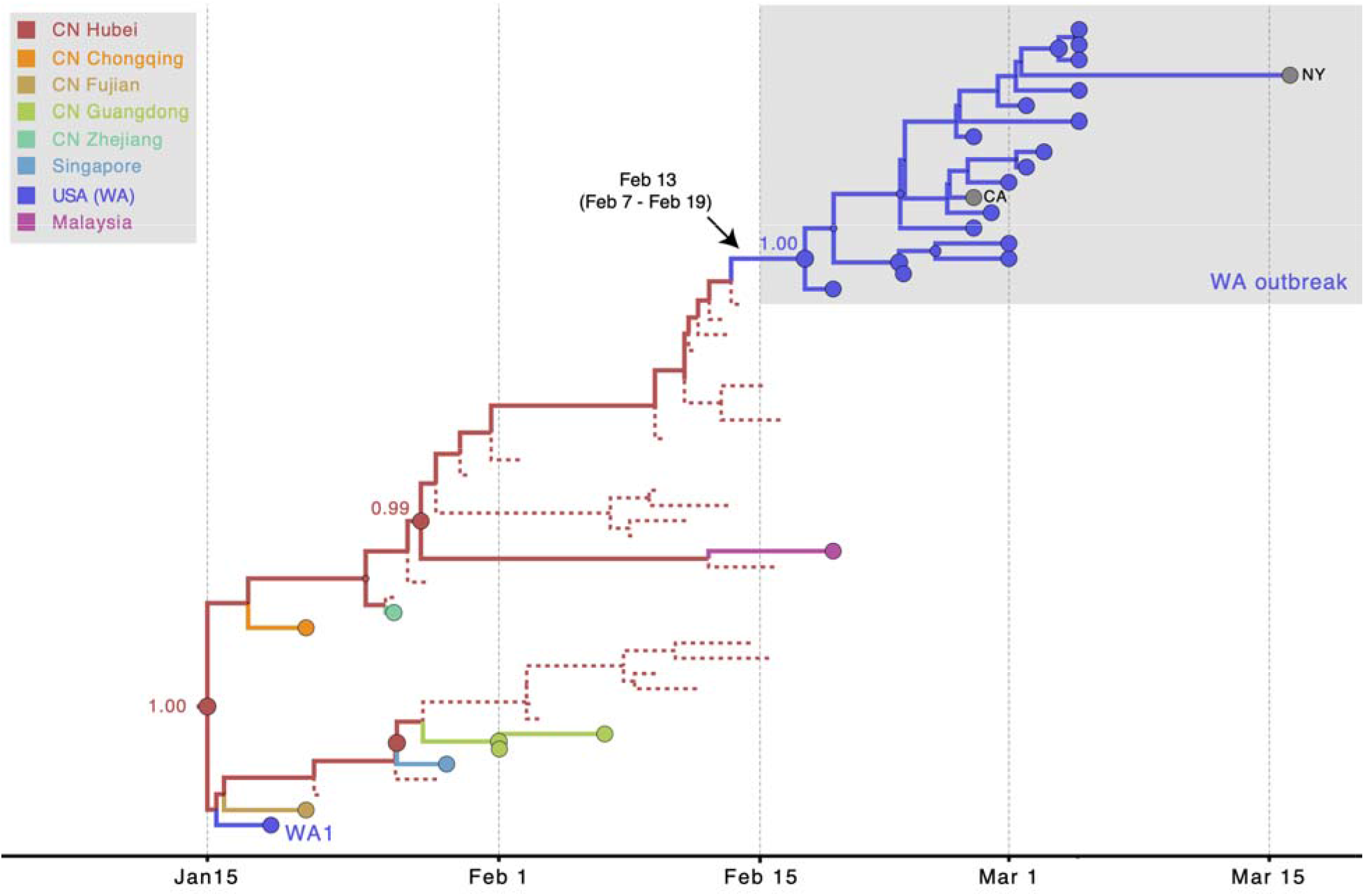
MCC tree of SARS-CoV-2 entry into Washington State. A subtree of the maximum clade credibility (MCC) tree depicting the evolutionary relationships inferred between (i) the first identified SARS-CoV-2 case in the US (WA1); (ii) the clade associated with the Washington State outbreak (including WA2); and (iii) closely related viruses that were identified in multiple locations in Asia. Circles at the tips represent observed taxa and are shaded by location. Branches and internal node circles are shaded similarly by posterior modal location state. Dotted lines represent branches associated with unsampled taxa assigned to Hubei, China (CN). Circle sizes for internal nodes are proportional to posterior clade support. Posterior location state probabilities are shown for three well-supported key nodes.

This timing is approximately four weeks later than had been proposed (*17*), implying: (a) archived ‘self-swab’ samples may have retrospectively detected the virus within as little as a week of its arrival (*5*), (b) the Washington State outbreak may have been smaller than estimated based on the earlier inferences, and (c) the individual who introduced the founding virus likely arrived in the US after the initiation of the ‘Suspension of Entry’ of non-US residents from China on February 2^nd^, 2020 (*20*) but during the period when an estimated 40,000 US residents were repatriated from China, with screening described as cursory or lax (*21*). These passengers were directed to a short list of airports including Los Angeles, San Francisco, New York, Chicago, Newark, Detroit and Seattle (*21*). So, although our reconstructions incorporating unsampled lineages do not account for travel restrictions, the remaining influx likely provided ample opportunity for a second introduction to Washington State. It is also possible that the virus entered via nearby Vancouver, British Columbia, which is closely linked to both China and Washington State.

### Early establishment of SARS-CoV-2 in Europe

Our simulation framework also suggested that the initial outbreak in Bavaria, Germany (represented by BavPat1) was unlikely to be responsible for seeding the Italian outbreak (see Fig. S1 for detailed phylogenetic scenarios). We simulated the origins of the Italian outbreak that was associated with viruses genetically related to BavPat1, again using realistic epidemiological parameters. Simulations with a median doubling time of 3.4 days (95% range: 2.9–4.4 days) resulted in a median epidemic size of 724.5 (95% range: 140–2,847) after 36 days. In the observed phylogeny, the Italian outbreak is the sole descendant lineage from BavPat1. Within the Italian outbreak, there are zero viruses identical to BavPat1 and four of the 27 related viruses included in this analysis are separated from BavPat1 by a single mutation. In simulation, the distributions of identical and one-mutation divergent viruses are not significantly different from expectation (*p*=0.146 and *p*=0.153, respectively). However, the lack of at least one descendent lineage that forms a polytomy with BavPat1 and the Italian outbreak significantly differs from expectation (*p*=0.005). Therefore, it is highly unlikely that BavPat1 or a virus identical to it seeded the Italian outbreak (Fig. S1).

An alternative scenario in which both the Germany and Italy outbreaks were independently introduced from China is further supported when missing sequences from undersampled locations are explicitly accommodated in the phylogeny. Using the approach described above, the evolutionary relationships of BavPat1 and viruses from the Italian outbreak were reconstructed while accommodating unsampled viruses assigned to Hubei and other locations (including Italy) determined to be undersampled, based on incidence data. The resulting phylogeny strongly supports independent viral introductions from Hubei into Germany and Italy, with the ancestral location states in Hubei supported by a posterior probability of 0.79 (Fig. 5). These findings again reveal that epidemiological linkages inferred from genetically similar SARS-CoV-2 viruses associated with outbreaks in different locations can be highly tenuous, given low levels of sampled viral genetic diversity and insufficient background data from key locations.

**Fig. 5.**
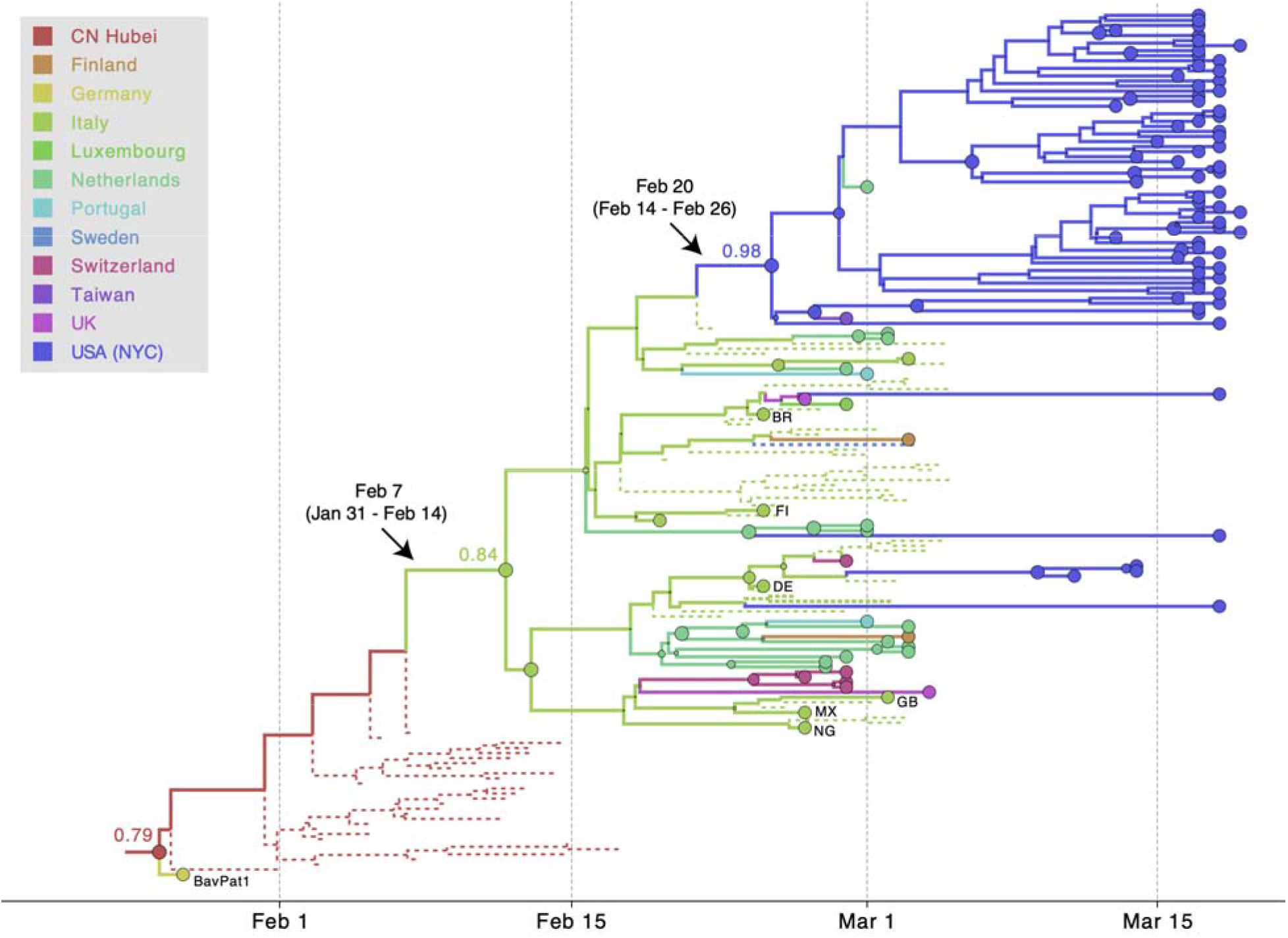
MCC tree of SARS-CoV-2 entry into Europe. A subtree inferred for viruses from (i) the first outbreak in Europe (Germany, BatPat1), (ii) outbreaks in Italy and New York, and (iii) other locations in Europe. Dotted lines represent branches associated with unsampled taxa assigned to Italy and Hubei, China (CN). Country codes are shown at tips for genomes sampled from travellers returning from Italy. Other features as described in Figure 4.

Interestingly, our approach is also able to infer that this major European clade, the same one that dominates in New York City (*14*, *15*) and Arizona (*22*), had an origin in Italy, as might be expected from the epidemiological evidence. This inference makes use of the recent travel history, where available, of various sequences in that cluster from people who had recently travelled to Italy. There are only two samples available from Italy in that cluster, yet we can trace strong evidence of the origin of this important lineage to Italy via Hubei. The Markov jump estimates of the movement from Hubei to Italy were: Feb. 7^th^, 2020 (95% HPD Jan. 31^st^–Feb. 14^th^). This Italian/European cluster, in turn, was the source of multiple introductions to New York City (NYC) (*14*, *15*). Using the same approach, we date the introduction leading to the largest NYC transmission cluster to Feb. 20^th^, 2020 (95% HPD Feb. 14^th^–Feb. 26^th^). Hence, even with its corrected age, the WA outbreak clade predates identified transmission clusters elsewhere in the US (*22*–*24*).

This global genomic perspective (Fig. 6) is relevant to recent claims (*16*) that this lineage is more transmissible due to the D614G Spike protein mutation. Instead, our results suggest the lineage possessing this mutation rode at least three early waves of uncontrolled transmission, first in the original Chinese epicenter of the outbreak, then in the earliest European epicenter in Northern Italy, and then in the uncontrolled outbreak in New York City. In other words, this viral lineage appears to have been amplified because of luck, not high fitness. We hypothesize that from these large centers of uncontrolled spread, where SARS-CoV-2 reached comparatively very high prevalence, D614G simply swamped D614 variants in other geographic regions (*22*, *25*). While we await definitive evidence, what is clear is that sensational claims of functional importance of this mutation were premature.

**Fig. 6.**
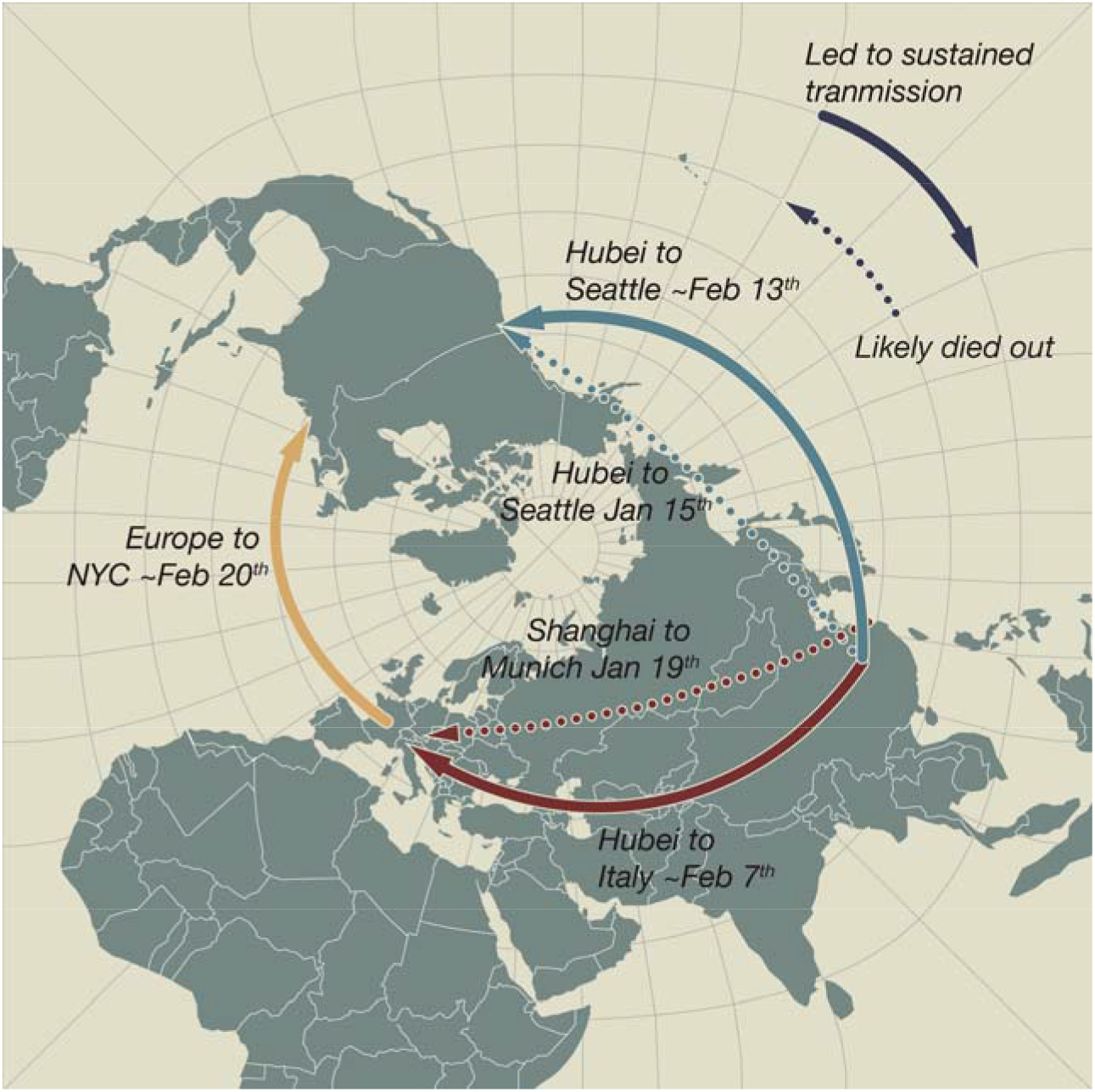
SARS-CoV-2 introductions to Europe and the US. Pierce projection mapping early and apparently ‘dead-end’ introductions of SARS-CoV-2 to Europe and the US (dashed arrows). These were followed by a series of dispersals (solid arrows) all likely taking place in February 2020: from Hubei Province, China to Northern Italy, from Hubei to Washington State, then from Europe (as the Italian outbreak spread more widely) to New York City.

## Discussion

January and February 2020 were pivotal months as government officials endeavored to understand and appropriately respond to the unfolding SARS-CoV-2 emergency, in the midst of considerable scientific uncertainty. When a new case was confirmed in a city that was not directly associated with travel, it was difficult to ascertain how the virus had gotten there and whether there had already been community transmission. There was also considerable uncertainty about whether epidemiological contact tracing and isolation would be effective for controlling new outbreaks. Weeks after the first diagnostic tests became available for SARS-CoV-2 on January 13^th^ (*26*), the story of German public health workers successfully using testing, contact tracing and isolation to control a small outbreak in Germany had profound implications for the feasibility of early, intensive interventions to prevent the virus from becoming established in Germany and other countries (*10*). Assertions that these efforts had actually failed may have led to confusion about the utility of these approaches and contributed to a sense of the inevitability of the spread of the pandemic. Similarly, conclusions that the Seattle area was already six weeks into an epidemic by the end of February, rather than two or three, and the notion that stringent efforts to prevent spread had failed in the WA1 case, may have influenced decision-making about how to respond to the outbreak, including whether such measures were worth the effort.

Despite the early successes in containment, SARS-CoV-2 eventually took hold in both Europe and North America during February 2020: evidently first in Italy in early February, then in Washington State mid-February, and then in New York City later that month (Fig. 6). Our finding that the virus associated with the first known transmission network in the US did not enter the country until mid-February is sobering, since it demonstrates that the window of opportunity to block sustained transmission of the virus stretched all the way until that point. It is clear that early interventions can have outsized effects on the course of an outbreak, and the precise impact of the slow rollout of diagnostic tests in the US on the early stages of the pandemic is likely to be explored and debated for years to come, including the initially narrow criteria for who could be tested. Our findings critically inform such inquiries by delineating when community transmission was first established in the US and by providing clarity on the duration of the time window before SARS-CoV-2 establishment when contact tracing and isolation might have been most effective.

Our findings highlight the potential value of establishing intensive, community-level respiratory virus surveillance architectures, such as the Seattle Flu Study, during a pre-pandemic period. The value of detecting cases early, before they have bloomed into an outbreak, cannot be overstated in a pandemic situation (*27*). Given that every delay in case detection reduces the feasibility of containment, it is also worth assessing the impact of lengthy delays in FDA approval of testing the Seattle Flu Study’s stored samples for SARS-CoV-2.

By delaying COVID-19 outbreaks by even a few weeks in the US and Europe, the public health response to the WA1 case in Washington State, and a particularly impressive response in Germany to a substantial outbreak, bought crucial time for their own cities, as well as other countries and cities, to prepare for the virus when it finally did arrive. Erroneously suggesting that WA1 introduced the earliest US outbreak of SARS-CoV-2 obscured the societal and public health benefits produced by an attentive, collaborative, and thoughtful patient willing to work with public health workers to prevent the spread of SARS-CoV-2. One irony is the beneficial impact the decision of government officials in Washington State, to be among the first in the US to initiate restrictions on social distancing and size of gatherings, had even though the decision was founded at least in part on an assumption about the timing of community transmission not supported by the phylogenetic data (i.e. the belief that cryptic transmission had been ongoing since mid-January). This action may have closed the gap between the onset of sustained community transmission and mitigation measures in Washington State, compared to other locales like New York City, in ways that deserve careful reevaluation.

Because the SARS-CoV-2 evolutionary rate is slower than its transmission rate, many identical genomes are rapidly spreading. This genetic similarity places limitations on some inferences such as calculating the ratio of imported cases to local transmissions in a given area. Yet we have shown that, precisely because of this slow rate, when as little as one mutation separates viruses, this difference can provide enough information for hypothesis testing when appropriate methods are employed. Bearing this in mind will put us in a better position to understand SARS-CoV-2 in the coming years.

## Acknowledgements

We thank the patients and healthcare workers who made the collection of this global viral data set possible and all those who made viral genomic data available for analysis. We thank Niema Moshiri for his guidance on FAVITES.

## Authors contributions

Conceptualization:MW

Methodology:MW,JP,MAS,PL,JOW

Software:JP,MAS,PL,JOW

Validation:JP,MAS,PL

Formal analysis:MW,JP,PL,MAS

Investigation:MW,JP,BBL,JBJ,AR,MIN,VH

Resources:MW,PL,MAS

Data Curation: BBL,JBJ,VH

Writing - original draft preparation:MW,MIN

Writing - review and editing:MW,BBL,MAS,JOW,JBJ,AR

Visualization: BBL,JOW,AR

Supervision:MW,JOW

Project administration:MW

Funding acquisition:MW,MAS,JOW

## Data and materials availability

BEAST .xml file example, FAVITES simulated phylogenies, and the GISAID accession numbers for all sequences used in the analysis are hosted at https://github.com/Worobeylab/SC2_outbreak

## Supplementary Materials

Methods

Supplementary Text

Table S1

Figure S1

## Supplementary Materials

### Materials and Methods

#### Methods

##### Epidemic Simulation using FAVITES

We performed a series of epidemic simulations using FAVITES (Main Text Figure 2). First, we simulated contact networks in FAVITES (Main Text Figure 2.1a). The reported number of contacts per day varies, with Mossong et al. reporting a mean of 13.4 contacts per day across 8 European countries, with Italy having 19.8 contacts per day (*28*). We selected an intermediate value of 16 contacts per day (i.e., mean degree) within a 20,000 node contact network generated via a Barabási-Albert model.

Concurrently, we performed a forward simulation SIR (Susceptible-Infected-Recovered) model to generate a transmission network over this contact network (Main Text Figure 2.1b). Starting with a single seed among our 20,000 susceptible individuals, we applied an infectiousness parameter of 2.7 and a recovery time of 20 days. The SIR model and contact network are linked to document the propagation of infection through the network, creating the transmission network (Main Text Figure 2.2). A single viral lineage from each infected individual was sampled at a random point during their infection time to represent viral genotype sampling. The viral phylogeny in units of time (years) was sampled under a coalescent model using the Virus Tree Simulator package embedded in FAVITES. The final output utilized from FAVITES is the viral time-based phylogeny (Main Text Figure 2.3). All inputs for FAVITES can be found in Table S1 and the JSON input files are available at https://github.com/Worobeylab/SC2_outbreak.

We constrained our simulated epidemics to have a median doubling time of 4.7 days (95% range: 4.2-5.1), corresponding to prior phylodynamic analysis (*17*). All resulting time trees were subsampled to match the dates corresponding to the 294 sampling dates in the WA outbreak (Main Text Figure 2.4). If 294 precisely matched dates were not found in the phylogeny, we subsampled tips that were closest to the desired dates by expanding the range to a 3-day window. If there were not enough viruses to properly capture the sampling dates for the WA outbreak clade using this method, the simulation was excluded. We produced 1000 simulations for each set of data.

##### Sequence Simulation and Phylogenetic Inference

We simulated genetic sequences over the sub-sampled epidemic trees using Pyvolve (*29*). The simulations were restricted to 262 variant sites found in an alignment (*17*) of length 29,850 with 295 taxa comprising WA1 and the Washington outbreak clade. We applied a genome-wide evolutionary rate of 0.8×10^−3^ substitutions/site/year to the variant sites, which translates to 0.1244 substitutions/site/year across the variant sites: (0.8×10^−3^ × 29,850) / 262 (Main Text Figure 2.5). To parameterize the sequence evolution, we first inferred a maximum likelihood tree for the 295 taxa WA outbreak clade alignment using a GTR + Inv model in IQ-TREE2 (*30*). We then applied the GTR substitution model to evolve the 262 variant sides down our simulated, sub-sampled epidemics, starting each simulation with the WA1 seed sequence (Main Text Figure 2.6). We then added the WA1 seed sequence along with the invariant genomic positions. A maximum likelihood phylogeny was then inferred in IQ-TREE2 using a GTR + Inv substitution model (Main Text Figure 2.7,8).

##### Sensitivity Analysis—Slower rate of infection

The same methods for epidemic and sequence simulation and tree inference were done on the WA data to test a more slowly spreading virus. We used the aforementioned parameters, except the contact network was 5,000 nodes and the infectiousness rate was 2.0-2.2 (Table S1). We constrained our output to be between 500 and 2,000 infected people and produced 1,000 replicates. The same methods as earlier were used for the subsequent tree pruning and scaling, sequence evolution, and tree inference.

##### Sensitivity Analysis—Faster rate of evolution

We took 500 of the subsampled mutation-scaled trees from the original WA epidemic and sequence simulation and multiplied each branch length by 2 to simulate an accelerated mutation rate of 1.6×10^−3^ substitutions/site/year. The same methods as earlier were used for the subsequent sequence evolution and tree inference.

##### Italy Outbreak Epidemic Simulation

The same methods for epidemic and sequence simulation and tree inference were used to model the Italy outbreak. We used a contact network of 10,000 nodes and the infectiousness rate was 2.6-2.8 for the epidemic simulation within FAVITES: 500 applied an infectiousness rate of 2.6, and the other 500 applied an infectiousness rate of 2.8. We constrained our simulated epidemics to have a median doubling time of 3.4 days (95% range: 2.9-4.4 days), similar to prior epidemiological analysis (*10*). The subsampling method remained the same. The Pyvolve simulations were restricted to 45 variant sites found in an alignment of length 29,775 with 28 taxa comprising Italian outbreak. The evolutionary rate of 0.8×10^−3^ substitutions/site/year was translated to 0.529 substitutions/site/year across the 45 variant sites: (0.8×10^−3^ × 29,775) / 45. The remainder of the pipeline was the same, except using the Germany/Italy data and seeding the evolution with the BavPat1 sequence. The files used for FAVITES, the viral time trees, and the subsampled mutation trees can be found in https://github.com/Worobeylab/SC2_outbreak.

##### SARS-CoV-2 genome data set

To investigate the evolutionary origins of early outbreaks of the SARS-CoV-2 virus in the Europe and US, we compiled a curated SARS-CoV-2 genome sequence data based on available genomes from the Global Initiative on Sharing All Influenza Data’s (GISAID) on March 10^th^, 2020 and genomes from New York City that subsequently became available with a collection date up to March 19^th^. Acknowledgements of all laboratories that contributed SARS-CoV-2 sequence data used in our study are found in https://github.com/Worobeylab/SC2_outbreak. The genomes were aligned using MAFFT v.7 and partially trimmed at the 5’ and 3’ ends. Viruses with evidence of sequencing error were removed and only a single genome was retained from patients for which multiple genomes were available. For additional quality control, root-to-tip divergence was visualized as a function of sampling time using TempEst (*31*), based on a maximum likelihood (ML) tree inferred with IQTree (*30*). One potential outlier was identified and removed. Genomes sampled from cruise ship travelers were not considered. The final data set consisted of 364 genomes that were sampled from 28 countries. Chinese samples were available from 13 provinces, one municipality (Beijing), and one special administrative area (Hong Kong), which were considered as separate locations in our phylogeographic reconstructions. The exact date of virus collection was available for all sequences except for one genome from Anhui, China, and 17 from Shandong, China. For viruses for which only the month of viral collection was available, the lack of tip date precision was accommodated by sampling uniformly across a 30-day window (*32*).

##### Bayesian phylogeographic inference

The evolutionary history of SARS-CoV-2 was inferred using a Bayesian approach, implemented through the Markov chain Monte Carlo (MCMC) framework available in BEAST 1.10.4 (*33*), using the BEAGLE library v3 (*34*) to increase computational performance. We specify an HKY85 nucleotide substitution model using a continuous-time Markov chain (CTMC) process, with a proportion of invariant sites and gamma-distributed rate variation among sites. An uncorrelated relaxed molecular clock model with a lognormal distribution was used with an exponential growth coalescent tree prior. The relatively low sequence variability over the short time scale of viral dissemination introduces considerable uncertainty into phylogenetic reconstructions of SARS-CoV-2.

Uncertainty in inferences of viral spatial movements is further compounded by the low availability of genetic sequence data from a number of critical countries and regions. To address such limitations, Bayesian reconstructions of the evolutionary history and spatial origins of SARS-CoV-2 were augmented by incorporating various sources of information. For genomes from patients with a recent travel history, we used the travel locations in our ancestral location reconstructions (except for WA1). For locations that are poorly represented (e.g. Italy), genomes sampled from travelers upon returning from these locations assist in uncovering the diversity in these locations.

##### Incorporating unsampled viruses into Bayesian phylogeographic inference

For poorly sampled locations, we included in the Bayesian inference unsampled genomes that were associated with an epidemiologically-informed sampling time and location but not observed sequence data. Phylogeographic reconstructions are highly sensitive to sample bias, and 13 locations were identified as critically under-sampled by contrasting the number of available genomes against the cumulative number of recorded COVID-19 cases in each location on March 10^th^, 2020. A standard approach to reduce geographical bias would be to down-sample data sets by removing genomes from more densely sampled locations. However, sequence data are sparse during the pre-pandemic stages of SARS-CoV-2 emergence and global spread, particularly from locations that experienced early outbreaks. In order to retain all available sequences, we opted instead to add unsampled taxa from underrepresented locations until achieving an arbitrary minimal ratio of 0.005 of taxa (sampled and unsampled) to cumulative number of cases in each location. To meet this threshold, 458 unsampled taxa were included from 13 locations that experienced early SARS-CoV-2 outbreaks but were poorly represented in the data available in GISAID. The majority of unsampled taxa were assigned to Hubei (*n* = 307), followed by Italy (*n* = 47), Iran (*n* = 40), and South Korea (*n* = 30).

Epidemiological information also was utilized to specify realistic sampling times for the 458 unsampled taxa. For each of the 13 locations associated with unsampled taxa, the time of sampling was described by a probability distribution, the shape of which was based on prevalent infections over time, estimated using the methods (and data sources) recently described by Fauver et al. (*24*). Probabilistic prior distributions were normally distributed (and appropriately truncated) in east Asian locations that effectively controlled early SARS-CoV-2 outbreaks through intensive social distancing and contact tracing. Probabilistic prior distributions were exponentially distributed in European countries where SARS-CoV-2 cases were still increasing exponentially in early March. Inaccuracies in COVID-19 case counts arising from underreporting and low testing are likely to affect the absolute number of COVID-19 cases in a given location, but less likely to impact the general shape and timing of epidemic curves, as used here. The probabilistic distributions for the sampling times of unsampled taxa were integrated over all possible times using MCMC in the full Bayesian analysis.

##### Bayesian phylogeographic inference with covariates

The final discrete spatial diffusion model included 44 locations, including 13 locations associated with unsampled taxa. To avoid an exceedingly high number of location transition parameters in a high dimensional CTMC, and to further inform the phylogenetic position of unsampled taxa, we utilized a generalized linear model (GLM) formulation of the discrete trait CTMC that parametrizes transition rates as a function of potential covariates (*35*). We considered three covariates related to human mobility that potentially predict long-distance routes of early SARS-CoV-2 spread: (i) air travel [as previously used in (*35*)], (ii) geographic distance (within continents), and (iii) an estimable asymmetry coefficient for Hubei, since in the early stage SARS-CoV-2 spread is dominated by an asymmetric flow out of Hubei, where the pandemic originated (*36*). The effect size of each covariate was estimated, along with an inclusion probability. In our phylogeographic inference, we also included a Markov jump counting procedure in order to estimate the times of specific transitions between locations (*19*, *35*). The posterior distribution of the full probabilistic model using MCMC sampling was estimated after running chains sufficiently long to ensure adequate effective sample sizes for continuous parameters, as diagnosed using Tracer (*37*). Posterior tree distributions were summarized using maximum clade credibility (MCC) trees that were visualized in FigTree.

### Supplementary Text

#### Detection of WA outbreak sequences

To determine the time frame that sequences identical to the WA outbreak clade putative common ancestor were sampled, we downloaded an aligned, complete genome dataset provided by GISAID (accessed May 11th, 2020) and inferred a neighbor joining (NJ) phylogeny with the HKY substitution model in Geneious Prime 2019.1.3. Sequences that appeared to be identical in the phylogeny to the WA outbreak putative common ancestor were extracted in Geneious and manually examined to make sure they did not contain an ‘N’ at any site that would make them non-identical. The earliest and latest genomes containing the key mutations C17747T and A17858G were sampled at the earliest on Feb 20^th^ (EPI_ISL_413456) and at the latest on April 27th (EPI_ISL_444051). This implies sequences identical to the WA outbreak clade putative common ancestor were circulating and detected for a minimum of 67 days. In contrast, the WA1 sequence was not detected again in another patient in Washington, despite intensive sampling. We do note however, that a single sequence identical to WA1 was sampled on March 13^th^, 2020 in NYC (EPI_ISL_427531).

#### Simulating a slower epidemic doubling time in Washington State

We simulated 1000 epidemics seeded by WA1 on January 15^th^ 2020 with a slower median doubling time of 5.6 days (95% range: 5.2–5.9) and the original evolutionary rate of 0.8×10-3 substitutions/site/year. These simulated epidemics produced a median of 938 cases after 61 days (95% range: 681–1,779 cases). Of the 1000 simulations, 500 applied an infectiousness rate of 2.0, and the other 500 applied an infectiousness rate of 2.2. As in the origins simulations, we observed a median distance from WA1 to the subsampled viruses of 3 mutations. In 77.8% of simulations, we observed at least one identical virus to WA1, with a median of 9 identical viruses in each simulation (95% range: 0–59 identical viruses). Not observing a viral sequence identical to WA1 in the real Washington data does not significantly differ from expectation (*p*=0.222). However, viruses with one mutation from WA1 were observed in 96.8% of simulations, indicating a low probability of not detecting a single sequence from Washington within one mutation of WA1 in the real data (*p*=0.032). Lineages forming a basal polytomy with WA1 and the epidemic clade were observed in 99.7% of populations (*p*=0.003) and 100% of simulations had at least one sibling lineage diverging prior to the formation of the outbreak clade (*p*<0.001). Therefore, our results are robust to an assumption of slower epidemic growth and smaller size.

#### Simulating a faster rate of evolution in Washington State

To explore the influence of a faster evolutionary rate on our simulations, we selected a random sample of 500 time trees from the main WA outbreak simulations and evolved the sequences with a substitution rate of 1.6×10^−3^ substitutions/site/year. These 500 simulated epidemics produced a median doubling time of 4.7 days (95% range: 4.2–5.1) and a median of 4204.5 cases after 61 days (95% range: 1996.5–11249.5 cases). In a departure from the true phylogeny and our previous simulations, the median distance from WA1 to the subsampled viruses evolved under a faster evolutionary rate was 6 mutations. In 29.2% of simulations, we observed at least one identical virus to WA1, with a median of 0 identical viruses in each simulation (95% range: 0-16 identical viruses). Not observing a virus identical to WA1 in the real Washington data does not significantly differ from expectation (*p*=0.708). In this case, viruses with one mutation from WA1 also did not significantly differ from expectation either (*p*=0.330). However, lineages forming a basal polytomy with WA1 and the epidemic clade were observed in 98.6% of populations (*p*=0.014) and 100% of simulations had at least one sibling lineage diverging prior to the formation of the outbreak clade (*p*<0.001). In sum, the faster evolution scenario produced trees that experienced more evolutionary divergence than observed in real-data, we still find strong evidence against WA1 serving as the founder of WA outbreak clade, as all of the simulated epidemics experienced a divergent lineage prior to the formation of the outbreak clade.

**Table S1.**
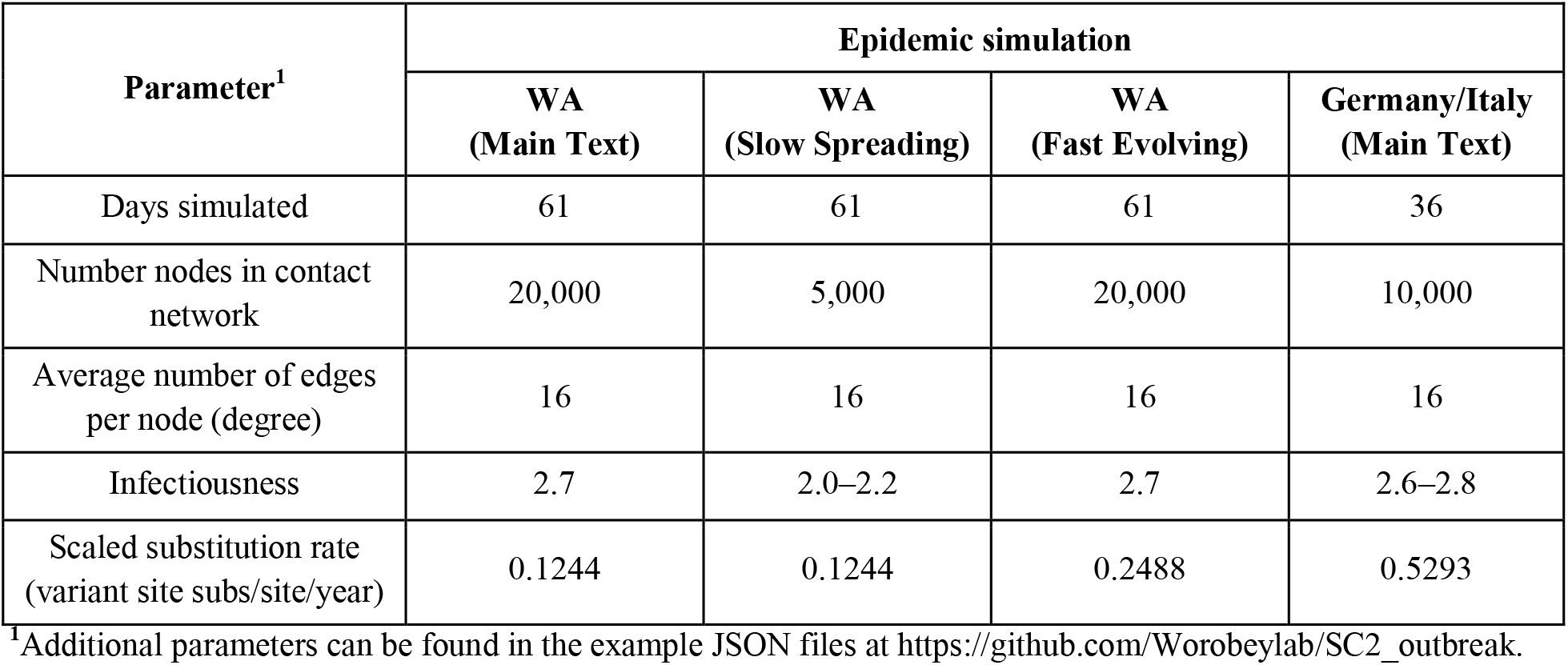
Parameters for epidemic and sequence simulations.

**Figure S1.**
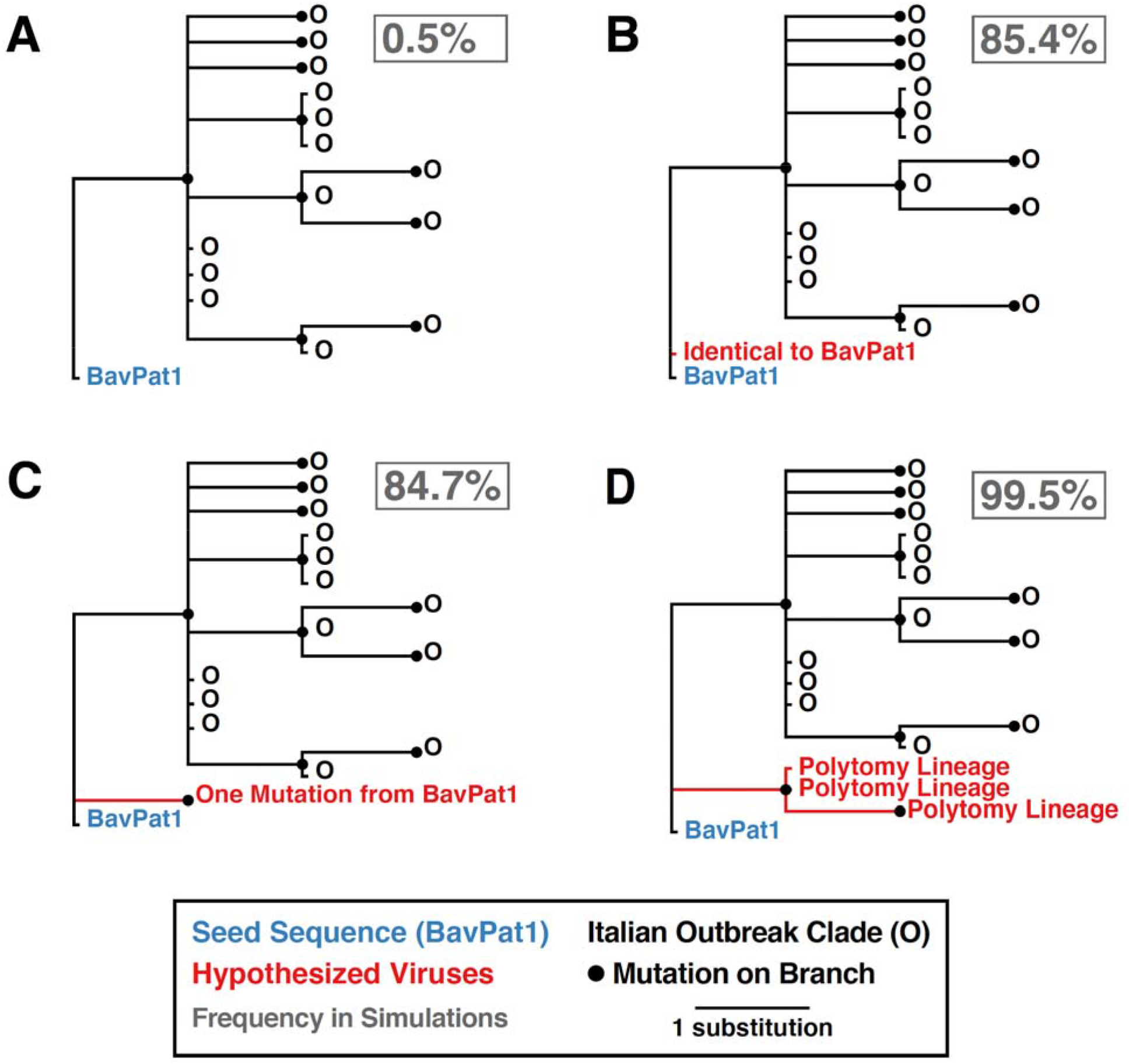
Potential phylogenetic relationships between BavPat1 and the Italian outbreak and their occurrence frequencies in 1000 epidemic simulations. (A) Observed pattern where the BavPat1 genome is the direct ancestor of the outbreak clade, separated by at least one mutation. (B) Identical sequence to BavPat1. (C) Sequences that are one mutation divergent from BavPat1. (D) Lineage forming a basal polytomy with BavPat1 and the outbreak clade. Frequency each relationship was observed in 1000 simulations reported in gray box.

